# Improving Social Interactive Learning through Dual Brain Stimulation

**DOI:** 10.1101/762377

**Authors:** Yafeng Pan, Giacomo Novembre, Bei Song, Yi Zhu, Yi Hu

## Abstract

Social interactive learning denotes the ability to acquire new information from a conspecific – a prerequisite for cultural evolution and survival. As inspired by recent neurophysiological research, here we tested whether social interactive learning can be augmented by exogenously synchronizing oscillatory brain activity across an instructor and a learner engaged in a naturalistic song-learning task. We used a dual brain stimulation protocol entailing the trans-cranial delivery of synchronized electric currents in two individuals simultaneously. When we stimulated inferior frontal brain regions, with 6 Hz alternating currents being in-phase between the instructor and the learner, the dyad exhibited spontaneous and synchronized body movement. Remarkably, this stimulation also led to enhanced learning performance. A mediation analysis further disclosed that interpersonal movement synchrony acted as a partial mediator of the effect of dual brain stimulation on learning performance, i.e. possibly facilitating the effect of dual brain stimulation on learning. Our results provide a causal demonstration that inter-brain synchrony is a sufficient condition to improve real-time information transfer between pairs of individuals.

**Significance:** The study of social behavior, including but not limited to social learning, is undergoing a paradigm shift moving from single- to multi-person brain research. Yet, nearly all evidence in this area is purely correlational: inter-dependencies between brains’ signals are used to predict success in social behavior. For instance, inter-brain synchrony has been shown to be associated with successful communication, cooperation, and joint attention. Here we took a radically different approach. We stimulated two brains simultaneously, hence manipulating inter-brain synchrony, and measured the resulting effect upon behavior in the context of a social learning task. We report that frequency- and phase-specific dual brain stimulation can lead to the emergence of spontaneous synchronized body movement between an instructor and a learner. Remarkably, this can also augment learning performance.

## 1. Introduction

Learning through interactions with others is one of the most extraordinary skills of humans among other social species (1, 2). Learning new information from a conspecific is often indispensable for survival. Yet, the scientific study of social interactive learning, and its underlying neurophysiological processes, has begun only recently (3–5).

A fundamental prerequisite of social interactive learning is the presence of (at least) two individuals: one teaching something to another. Accordingly, the most recent brain research in this area is moving towards paradigms entailing the simultaneous recording of two individuals’ neural activity, and the analysis of their inter-dependency (6). This is also referred to as “hyperscanning” (7, 8).

In a recent hyperscanning study, we examined brain activity from dyads composed of instructors and learners engaged in the acquisition of a (music) song (4). We observed that neural activity recoded over the inferior frontal cortices (IFC) of the instructor and the learner become synchronized, particularly when the learner was observing the instructor’s behavior. Remarkably, inter-brain synchrony (IBS) predicted learning performance, in particular the learner’s accuracy in pitch performance learning (i.e. intonation).

Our observations join others in suggesting that IBS is a correlate of social interactive learning (3–5, 9). Yet, the functional significance of this phenomenon remains elusive. One could claim that synchronous brain activities occur as a consequence of social interactive learning. Alternatively, a stronger claim could suggest that IBS is a sufficient condition to enhance social interactive learning. If this was the case, then it should be predicted that exogenously enhancing IBS would cause improved learning performance.

To test the above hypothesis, we adopted a “dual brain stimulation” protocol (10). This consists of simultaneous electric currents delivered trans-cranially in two individuals simultaneously. By manipulating the coupling between the signals delivered across two brains, the experimenters can control IBS and monitor its causal effects upon social behavior (10).

Following up on our previous study (4), we targeted the IFCs of dyads composed of an instructor and a learner – engaged in the acquisition of a song – using pairs of transcranial alternating current stimulators (tACS). We delivered alternating currents oscillating in the theta frequency range (6 Hz) because oscillations in this band are commonly observed over the frontal cortex, specifically in the context of tasks requiring auditory processing (13, 14), musical interaction (11, 12), or learning (15, 16) – all prerequisites to our song learning task. Crucially, we manipulated the relative phase of the learner’s and the instructor’s currents (**Fig. 1**, *lower panels*), being these either perfectly in-phase (0° relative phase) or in anti-phase (180° relative phase). For control purposes, we also included a control stimulation frequency of 10 Hz, as representative of the alpha band (17), and a sham stimulation condition.

**Fig. 1.**
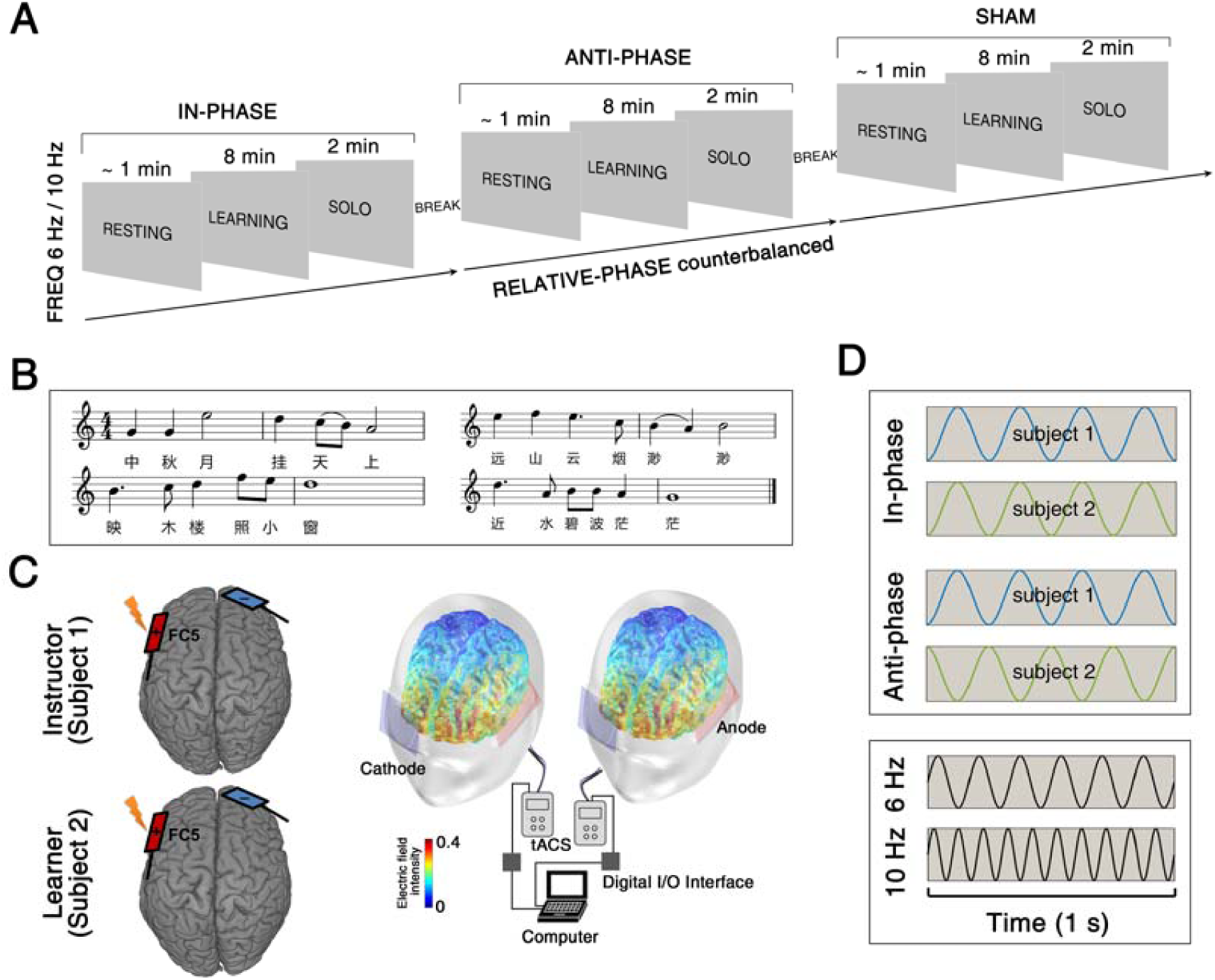
Experimental task and design. (A) The social interactive learning task consisted of three sessions: Resting, Learning, and Solo. (B) Representative musical score used in the social interactive learning task. (C) Schematic illustration of the brain stimulation montage. Left: dual brain stimulation was administered through simultaneous tACS to the instructor and the learner. The electrodes were placed over FC5 (anode) and FP2 (cathode), according to international 10/10 system, in order to best target the left inferior frontal cortices. Right: Electric field simulation shows that this montage entrains neural activity in the brain region of interest. (D) The dual brain stimulation protocol entails manipulations of relative-phase (between instructor and learner) in different frequencies.

We collected two distinct measures of behavior. On the one hand, we asked a group of expert raters to evaluate how well had the learners acquired the musical material. In line with our previous work, we expected 6 Hz in-phase dual brain stimulation to enhance intonation learning performance (4, 10). On the other hand, we used HD video recordings of the learning task to extract indices of spontaneous movement of the two participants. This second measure was meant to be exploratory. It was inspired by a growing body of research indicating that interpersonal synchronous movement can augment pro-social behaviors (18–23), hypothetically even social learning.

## 2. Results

The social learning task comprised two main sessions: a Learning session and a Solo session. During the Learning session, the instructor taught the song to the learner while their brains were simultaneously stimulated. Next, during the Solo session, learners were instructed to sing the newly acquired song as best as they could, while the performance was recorded in order to be evaluated by expert raters afterwards (**Fig. 1**). For consistency with the temporal order of the two sessions, we firstly present the results of the interpersonal movement synchrony analysis (associated with the Learning session) and later we report the experts’ ratings of the solo performances. Finally, we report correlation and mediation analyses addressing the relationship between interpersonal movement synchrony and learning performance.

### 2.1. Dual brain stimulation enhanced interpersonal movement synchrony

The results from the cross-correlation analysis are displayed on **Fig. 2 (*C*** and ***D***). The cluster-based permutation test on these data yielded evidence in favor of a main effect of stimulation FREQUENCY (lags −0.76 ∼ +0.40 s, *P* < 0.01), a main effect of RELATIVE-PHASE (lags −0.24 ∼ +0.24 s, *P* < 0.01) and an interaction between these two (lags −0.44 ∼ +0.40 s, *P* < 0.05). These results indicated that interpersonal movement synchrony was generally higher when the two brains were stimulated (*i*) at 6 Hz (0.17 ± 0.05), as opposed to 10 Hz (0.14 ± 0.03), and (*ii*) in-phase (0.19 ± 0.07), as opposed to sham (0.13 ± 0.03). These two effects appeared to be additive, as supported by a significant interaction indicating that interpersonal movement synchrony was maximal in the 6 Hz in-phase condition (0.22 ± 0.06).

**Fig. 2.**
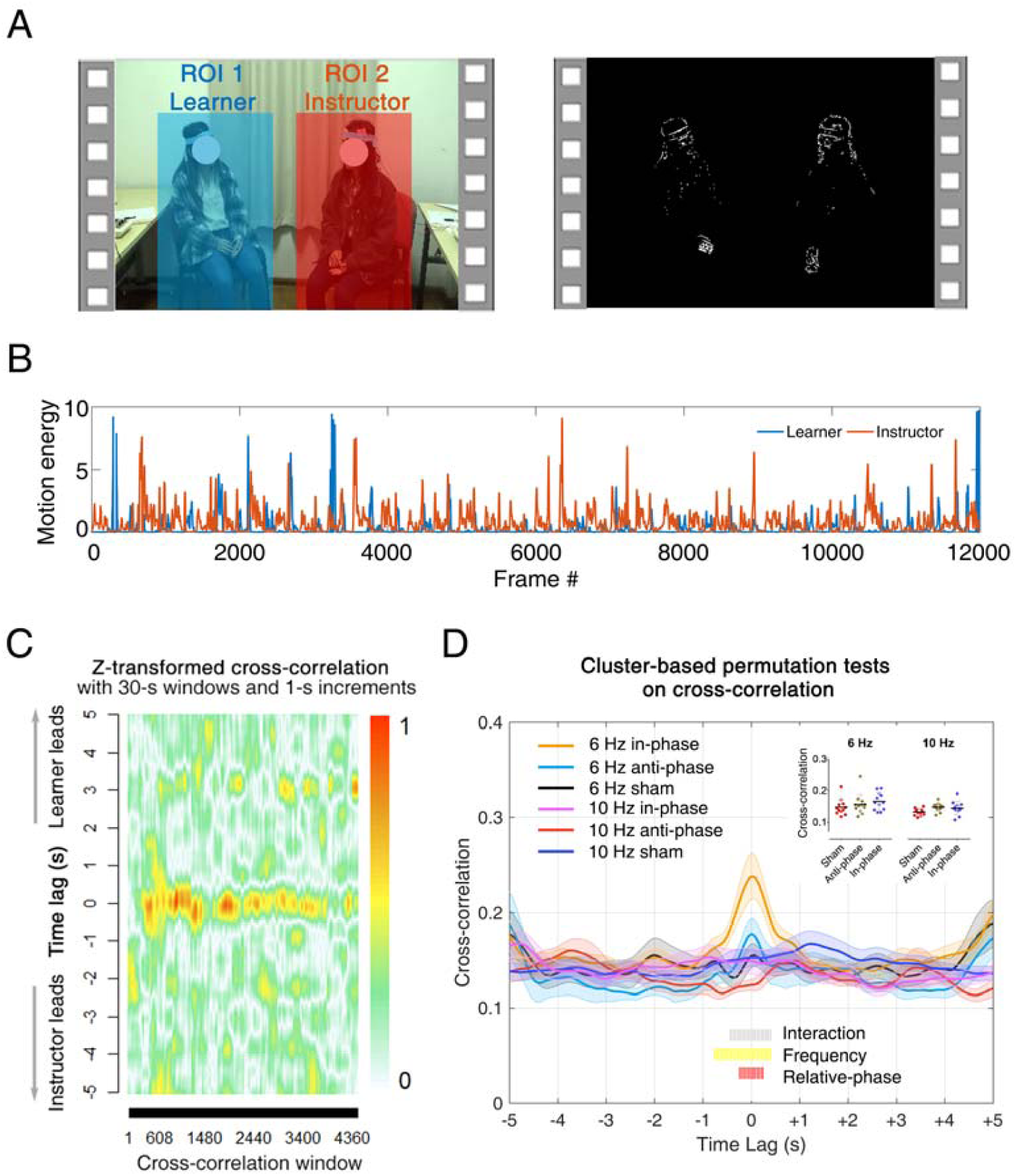
Motion energy extraction and cross-correlation analysis. (A) Regions of interest (ROI) utilized for the video-based Motion Energy Analysis. ROI1 covers the face and body of the learner (in blue); ROI2 covers the face and body of the instructor (in red). (B) Representative motion energy time series. (C) Cross-correlation coefficients obtained using a moving window approach (window size = 30 s; moving in steps of 1 s) in a representative dyad from the 6 Hz group. (D) Cross-correlation coefficients averaged across time, and participants, within each condition. The shaded area denotes the standard error at each time lag. We used a cluster-based permutation test to control for multiple comparisons. Periods of time associated with significant clusters are marked on the bottom. Single dyads’ coefficients (averaged within the interaction cluster) are plotted on the top-right side of the panel.

### 2.2. Dual brain stimulation improved intonation learning performance

Experts’ ratings of the solo performances (provided following the Learning session) are displayed on **Fig. 3**. Statistics indicated that intonation accuracy changed as a function of the dual brain stimulation condition. Specifically, the results yielded evidence for an interaction between RELATIVE-PHASE and FREQUENCY [*F*(2,44) = 6.86, *P* = 0.003, η^*2*^*partial* = 0.24]: raters judged intonation to be better for songs associated with in-phase stimulation (4.17 ± 1.68) compared to sham stimulation (3.46 ± 1.50, *P* < 0.05, Bonferroni corrected). Crucially, this was the case only for 6 Hz, but not 10 Hz, stimulation (*P*s > 0.05). The ANOVAs conducted on the ratings associated with the other musical aspects yielded no statistically significant effects, *F*s < 1.90, *P*s > 0.05 (**Fig. 3**).

**Fig. 3.**
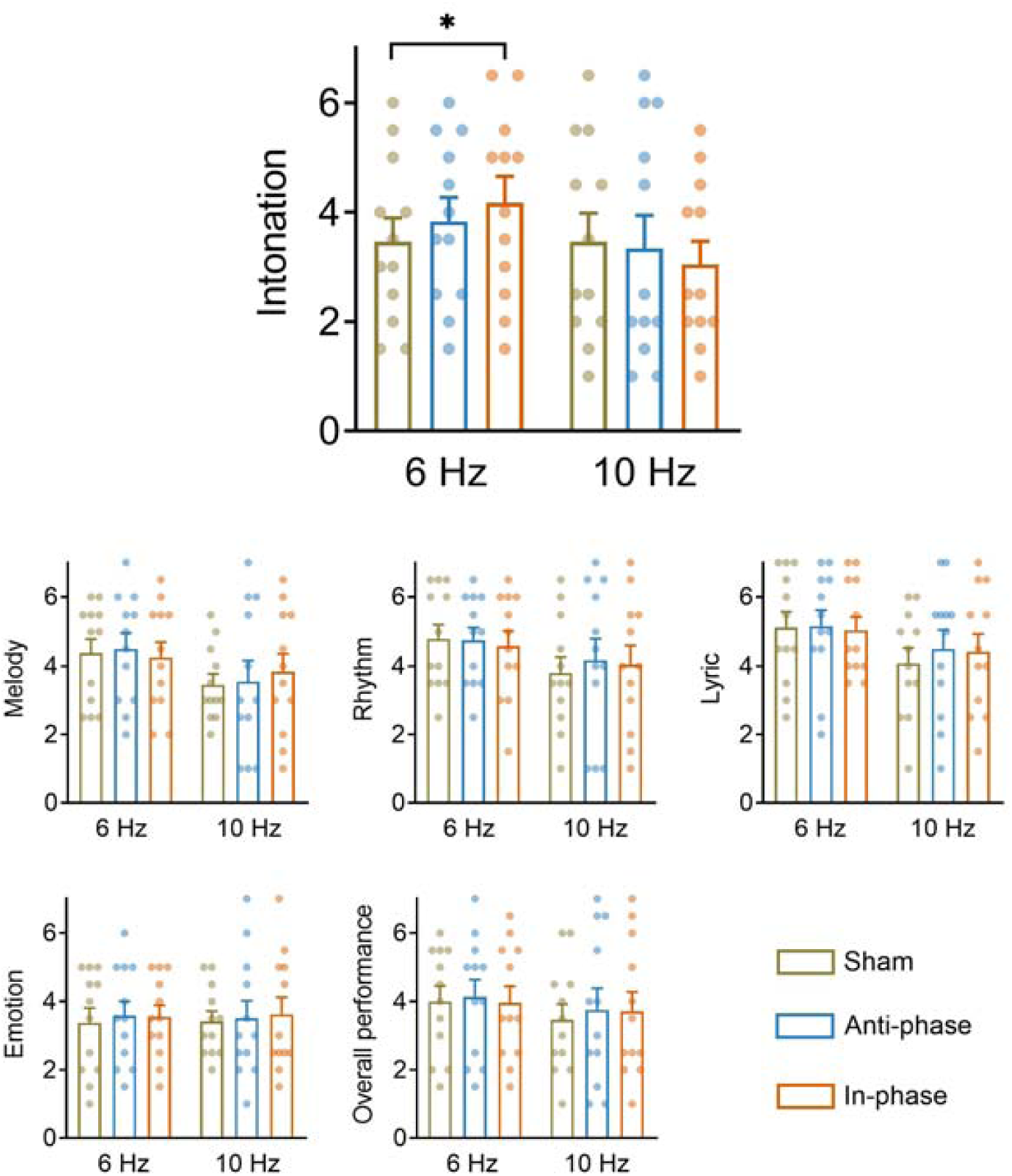
Learning performance ratings. Experts were asked to evaluate six aspects of learners’ solo performance (intonation, melody, rhythm, lyric, emotion, and overall performance – see Methods for details). Results revealed that 6 Hz in-phase dual brain stimulation led to better intonation compared to the sham stimulation. \**P* < 0.05. Error bar represents standard error. The other aspects of the performance were not affected by dual brain stimulation.

### 2.3. Interpersonal movement synchrony partially mediated the effect of dual brain stimulation on intonation learning performance

The results from the correlation analysis are shown on **Fig. 4*A***. The results indicated that the stimulation-mediated enhancements (referred to as “Δ”) of interpersonal movement synchrony and intonation learning performance were positively correlated (*r* = 0.62, *P* = 0.03). This implied that 6 Hz in-phase dual brain stimulation led to relatively more synchronized movement in those learners whose performance was also rated higher. This observation suggested that dual brain stimulation had possibly enhanced the acquisition of the musical material by inducing spontaneous and synchronized movement across the instructor and the learner.

**Fig. 4.**
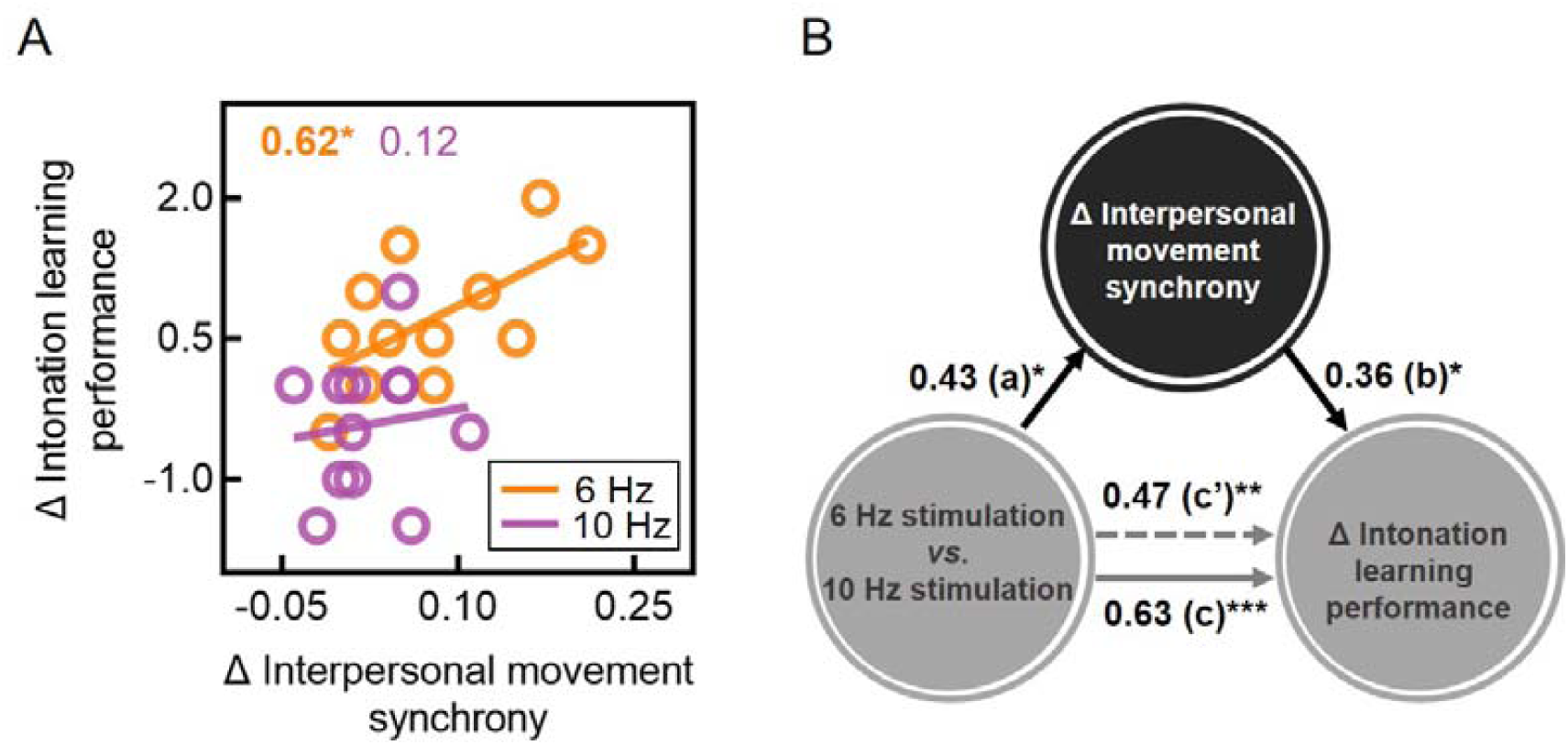
Correlation and mediation analyses. (A) Δ Interpersonal movement synchrony (in-phase minus sham) was positively correlated with Δ intonation learning performance in the 6 Hz (but not 10 Hz) group. (B) The effect of dual brain stimulation on Δ intonation learning performance (in-phase minus sham) is partially mediated by Δ interpersonal movement synchrony (in-phase minus sham). All path coefficients are standardized. \**P* < 0.05, \*\**P* < 0.01, \*\*\**P* < 0.001.

This hypothesis was confirmed by the results of the mediation analysis (**Fig. 4*B***). Notably, the results indicated (*i*) that dual brain stimulation could predict learning performance (path *c* = 0.63, *P* = 0.001), and (*ii*) that the relationship between dual brain stimulation and learning performance was reduced but still significant when interpersonal movement synchrony was included in the model as a mediator (path *c’* = 0.47, *P* = 0.004). This mediation effect was different from zero with 95% confidence (*β* = 0.26, confidence intervals = 0.01 to 0.74). Thus, interpersonal movement synchrony acted as a partial mediator of the effect of dual brain stimulation on intonation learning performance, possibly facilitating but not causing the effect of dual brain stimulation on learning.

## 3. Discussion

We report unprecedented evidence that dual brain stimulation can augment social interactive learning. Alternating currents were delivered simultaneously through the brains of learners and instructors – engaged in a social learning task entailing the acquisition of a song – targeting inferior frontal cortical (IFC) regions. When the exogenously controlled currents were programmed to both oscillate at 6 Hz, and with an in-phase relation across the learner and the instructor, we observed enhanced learners’ performance. Specifically, intonation learning performance following 6 Hz in-phase stimulation was rated as higher than following conditions implying 6 Hz anti-phase, sham stimulations or 10 Hz control conditions.

This result is particularly important because it fits nicely with our previous correlational evidence showing that – in the context of a similar song-learning task – the brains of learners and instructors synchronize, and the strength of such synchrony predicts learners’ intonation learning performance (4). Thus, going well beyond the previous observation, our current result indicates that inter-brain synchrony is not simply an epiphenomenon of social interactive learning, but a neurophysiological condition that per se is sufficient to enhance social interactive learning. This result speaks to a large community of scientists working in hyperscanning research addressing, besides learning, other topics such as cooperation (24), decision making (25), communication (26), and joint attention (27). It shows that the notions acquired using hyperscanning can be brought to a whole new frontier. Rather than seeking correlational evidence between inter-brain synchrony and social behavior, scientists could attempt to manipulate social behavior by controlling inter-brain synchrony. This could lead to a noteworthy paradigm-shift in social neuroscience, with remarkable applications touching on pedagogy, psychiatry, economics, and beyond (6, 28, 29).

Dual brain stimulation has been firstly developed by Novembre et al. (10), and later used by others (30). In their first study, Novembre et al. (10) targeted primary motor regions (M1) while two participants were preparing to tap their fingers in synchrony. It was reported that in-phase 20 Hz stimulation enhanced interpersonal coordination, specifically the dyad’s capacity to establish synchronized behavior. Based on our previous observations (4) and other music-related neuroscientific reports (31), our study was not meant to target M1, but rather IFC. For this reason, the stimulation was delivered more frontally (approximately over FC5 instead of C3 – see Methods). Furthermore, the delivered currents oscillated in the theta range (6 Hz), as opposed to the beta range (20 Hz), in accordance with relevant literature characterizing the neuroanatomical origin of such neural rhythms (32–34).

Nevertheless, because of the similar approach used in the two studies, we explored whether the two participants’ bodies swayed during the learning task, and whether those putative movements synchronized across learners and instructors. The reader should bear in mind that participants were not instructed to perform full-body movements. Thus, this analysis addressed spontaneous, as opposed to intentional (i.e. goal-directed) (35), body movement, which is meant to be a functionally distinct cognitive and neurobiological process (36, 37).

The analysis of interpersonal (spontaneous) movement synchronization yielded a very interesting result. Specifically, it showed how 6 Hz in-phase stimulation not only enhanced learning performance, but also led to enhanced movement synchronization between the learner and the instructor. Specifically, keeping in mind the structure of the experimental procedure, participants increased interpersonal movement synchronization while receiving 6 Hz in-phase dual brain stimulation and simultaneously learning the song (Learning session). Next, following this specific stimulation condition, intonation learning performance was found to be enhanced (Solo session).

This observation, and the temporal order of the effects, suggested that perhaps dual brain stimulation was not directly enhancing learning, but possibly it was doing so indirectly, i.e. through enhancement of interpersonal movement synchrony. This hypothesis was based on evidence indicating that interpersonal movement synchronization leads to noticeable pro-social effects such as enhanced partner likability, trust and affiliation (18–23) – all factors that might impact upon an interpersonal learning performance (9). To address this suggestion, we performed a mediation analysis. The results of this analysis, however, provided only partial support to this account. Specifically, we observed that interpersonal movement synchrony worked only as a partial mediator, and therefore it could explain the effect of brain stimulation on learning only in part.

It should be noted how the two proposed (not necessarily alternative) accounts of our results point towards markedly different underlying neurophysiological processes. The first account, according to which dual brain stimulation directly enhanced social learning, would be consistent with the broadly accepted notion that the phase of neural rhythms reflects periodic moments of enhanced cortical excitability (38, 39). When this phase, recorded from the brain of one individual, aligns with the phase of another individual, the pair benefits from a neural alignment that might improve information transfer and support lots of interpersonal activities (40). From this perspective, the instructor would have transferred information to the leaner more efficiently. Instead, the account based on pro-social behavior being driven by enhanced movement synchrony could call for other neural processes, such as social affective networks (41) or neurohormonal mechanisms regulating e.g. endorphins or oxytocin release (42, 43). Future research might attempt to shed light upon the role (and possible interplay) of these mechanisms in social interactive learning.

A few other outstanding questions remain unanswered. For instance, besides affecting intonation accuracy specifically, our previous study reported also an effect of IBS on overall learning. Why did our stimulation protocol affected intonation specifically, leaving overall performance unaffected? This difference could be explained by the different methodologies, and resulting timescales, used across our previous and current studies. Specifically, our IBS observations were made using fNIRS signals, which rely on hemodynamics and therefore unfold very slowly, resulting in ultra-low frequencies (below 1 Hz). Instead, the current study relying on tACS was designed taking into account electrophysiological neural rhythms, which are much faster and normally range in between 1 and 100 Hz. In this area, 6 Hz was selected as a promising rhythm due to its role in pitch processing (13) and auditory change detection (14). It follows that the current approach might have specifically targeted neural mechanisms responsible for intonation, while the previous measure of IBS might have captured additional ones.

A second point relates to sex composition of our cohort. Only female participants were tested in order to reduce variability of our sample, in accordance with previous evidence and recommendations (44–46). Although a strict criticism could question whether our results are generalizable to male individuals, we have no a priori reasons to expect so. Yet, being our effects sex-specific or sex-selective, we believe our results make a very important contribution to the emerging field of “multi-person neuroscience” (6).

## 4. Methods

### 4.1. Participants

We recruited twenty-eight healthy, right-handed volunteers: twenty-four of which acted as learners (mean age ± SD: 20.96 ± 2.31, age range: 17–25), while the remaining four acted as music instructors (mean age ± SD: 19.25 ± 0.50, age range: 19–20). We tested only female-female participant dyads in order to mitigate inter-individual and inter-dyad variability (44, 45), in accordance with recent work (4, 46). The four instructors were required to have received at least 10 years of formal musical training and they were all members of a local choir. The twenty-four learners were required to have (*i*) less than 3 years of formal musical training and (*ii*) no musical training at all within the past 5 years. Each of the four instructors was paired with 6 learners, in a one-by-one fashion, resulting in a total of 24 instructor-learner dyads. None of the participants had a history of neurological or psychiatric illness. All participants were naïve with respect to the purpose of the study. Each participant provided informed consent prior to the experiment and was paid for participation. The study was approved by the University Committee of Human Research Protection (HR 125-2018) from East China Normal University.

### 4.2. Experimental task

In a social interactive learning task, the instructor taught three songs to each learner individually while seating face-to-face (0.8 meters apart). The instructor and the learner’s chairs were slightly oriented towards the camera to improve whole-body visibility (resulting in approximately a 90° angle in between the two chairs’ orientations). Each song was taught within a dedicated block, which comprised three sessions: Resting, Learning, and Solo (**Fig. 1*A***). During the Resting session (∼1 min), the instructor and the learner were asked to relax and to avoid unnecessary movement. During the following Learning session (8 min), the instructor taught the song to the learner in a turn-taking manner, i.e. the learner attended and then imitated every single phrase of the song (i.e. one-by-one) performed by the instructor (4). Note that the learning task was meant to unfold in a naturalistic manner, and therefore both the instructor and the learner were free to use vocal and non-vocal communication (including facial expressions or gestures) to facilitate the acquisition of the song. Finally, during the Solo session (2 min), learners were instructed to sing the whole song as best as they could. This allowed us to record the final performance and later assess how well the song had been acquired.

### 4.3. Musical material

We selected three Chinese songs conveying a similar musical structure (e.g., quadruple rhythm, eight bars, and slow tempo) and emotion (i.e., missing home): (*i*) “The Moon Reflection” (Lyrics: B. Peng, Music: Z. Liu and S. Yan), (*ii*) “Nostalgia” (Lyrics: T. Dai, Music: Z. Xia), and (*iii*) “A Tune of Homesickness” (Lyrics: C. Qu, Music: Q. Zheng) (**Fig. 1*B*** displays a segment from “Nostalgia”). These songs were selected because they were meant to be unfamiliar to the learners (as confirmed by learners’ report), and because they were used in our previous work motivating the current study (4).

### 4.4. Experimental design

During (and only during) the Learning session, we delivered tACS to both the learner and the instructor using a dual brain stimulation protocol (see the next section for technical details). Our experimental design entailed two manipulations. First, we manipulated the FREQUENCY of the induced current, being 6 Hz for half of our participants, and 10 Hz for the remaining half. Note that age, prior musical training (in years), pitch discrimination and music memory abilities (as assessed by a 6-min online test, http://jakemandell.com/tonedeaf/) (47) were comparable across these two groups (*t*s < 0.79, *P*s > 0.44). Second, we manipulated the RELATIVE-PHASE of the signals delivered across the instructors and the learner. These could be either in-phase (i.e. 0° relative phase) or anti-phase (i.e. 180° relative phase) (**Fig. 1*D***). A third sham stimulation condition was also included for control purposes, leading to a full 2 × 3 factorial design (see *SI Appendix*, **Table S1**, for more details).

### 4.5. Dual brain stimulation

A dual brain stimulation protocol was used to deliver simultaneous signals to the brains of the instructor and the learner during the Learning session. To achieve this, we used two battery-driven tACS stimulators (Model: 2001; Soterix Medical Inc., New York, USA). The signals were delivered trans-cranially through two electrodes covered with rubber (5 × 5 cm; Soterix Medical Inc., New York, USA) and soaked in a saline solution (5 × 7 cm; Soterix Medical Inc., New York, USA). All stimulation electrodes were secured to the scalp with rubber head straps. For both participants, the anode electrode was placed over the left IFC (equivalent to electrode position FC5 according to the international 10/10 system), while the cathode electrode was placed over the contralateral frontopolar cortex (**Fig. 1*C***) (48, 49). An electric field simulation obtained using the COMETS Toolbox (version 2.0) (50) confirmed that this montage is appropriate to entrain neural activity in the left IFC **(Fig 1*C***).

The stimulation entailed a sinusoidal wave having a peak-to-peak amplitude of 1 mA, which ended the ramping up phase when the Learning session began and began the ramping down phase when the session ended. The frequencies and relative phases adopted are described in the previous section. For sham stimulation conditions, both subjects received a 30 s fade-in followed by a 30 s fade-out of stimulation.

The two stimulators were controlled through a National Instruments Data Acquisition Toolbox Support Package (NI-DAQmx), which was controlled using MATLAB (MathWorks Inc., Natick, MA) via two USB/Parallel 24-Bit Digital I/O Interfaces (Model: SD-MSTCPUA; Cortech Solutions Inc., North Carolina, USA). The latter was connected to a computer running the Data Acquisition Toolbox as well as to each stimulator. An external trigger was sent simultaneously from the computer to the Digital I/O Interfaces so that the two stimulators could begin the stimulation at the same time (**Fig. 1*C***).

Prior to the experiment, all participants were exposed to tACS for approximately 1 minute to ensure they were comfortable with the stimulation intensity. Both the instructor and the learner were naïve with respect to the RELATIVE-PHASE and FREQUENCY conditions applied (see *SI Appendix* for details). After each experimental block, participants were asked to report potential side-effects of the tACS. They filled in a questionnaire including the following items: pain, burning, heat, itchiness, pitching, metallic taste, fatigue, skin flush, the effect on performance or any other side-effect perceived. Reported side-effects were comparable across conditions (*F*s < 1.14, *P*s > 0.12).

The instructors, who were meant to teach to six different learners and therefore undergo the stimulation procedure on six different occasions, were allowed to a maximum of two teaching sessions per week (with at least three days in between).

### 4.6. Video and audio recordings

The whole procedure was video-recorded using a fixed digital camera (HDR-CX610E, Sony, Tokyo, Japan). Recording files were stored using a MTS format. The room illumination was controlled in order to be stable and support optimal shutter-speed and aperture. The distance between the camera and dyads was about 2 meters so that both participants’ full bodies and faces could be captured. Additionally, a digital voice recorder (ICD-PX470, Sony, Tokyo, Japan) was used to record the vocalizations. The voice recorder was placed nearby the participants (∼30 cm). The audio files were stored in WAV format. The high-quality video and audio recordings were subsequently used to quantify movement dynamics and evaluate the song learning performance.

### 4.7. Data analysis

Two main analyses were conducted. First, the video data collected during the Learning session was analyzed in order to quantify whether the learner and the instructor movements synchronized. Second, we analyzed both the video and the audio recordings associated with the Solo session. These recordings were presented to a group of expert raters (naïve with regards to the purpose of the study), who rated how well the materials had been acquired. All of these analyses were firstly conducted within experimental conditions, and the results were later compared statistically (see next section).

#### 4.7.1. Video analysis of the Learning session

Preprocessing of the video data was conducted using the Format Factory (version 4.1.0, PCFreetime Inc., Shanghai, China). The MTS files were first converted into the MP4 format (FPS 25; dimension 1920 × 1080). The converted data (25 frames/s) was segmented according to the trial structure, i.e. separate segments, either associated with the Learning session or with the Solo session, were extracted. The analysis of the segments associated with the Learning session consisted of the following four steps.

##### Step 1: Instructor and learner movement extraction

A Motion Energy Analysis (MEA) algorithm was used to compute a continuous measure of movement associated with either the instructor or the learner (51). This algorithm employs a frame-differencing approach, i.e. it quantifies the amount of change from one frame to the next, i.e. motion energy (52). The algorithm was applied to two separate regions of interest, each covering the full body of the instructor or the learner (**Fig. 2*A***).

##### Step 2: Preprocessing of time series

The motion energy signals resulting from the previous step were smoothed using a moving average window (span = 0.5 s). Next, outlying data points within the time series (i.e. values exceeding mean + 10 * STD of the time series) were removed (1.23 ± 0.21% of the whole data).

##### Step 3: Cross-correlation analysis

The preprocessed time series were subsequently submitted to a cross-correlation analysis, which was meant to quantify the dynamic synchrony between the instructor and the learner. Before entering the data into this analysis, we controlled whether the mean and standard deviation of the time series (indexing the amount of movement, and movement variability) were comparable across conditions (all *P*s < 0.18). Next, motion energies associated to the instructor and the learner were cross-correlated, separately for each condition, using a moving window (span = 30 s, maximum lag = 5 s, step = 0.04 s, leading to 125 steps) (53). Note that the moving window approach is appropriate considering the non-stationary nature of movement behaviors.

##### Step 4: Interpersonal movement synchrony

Cross-correlation coefficients comprised 251 time lags (125 cross-correlations for positive lags, 125 for negative lags, and 1 for the zero lag). These were Fisher’s *z* transformed to obtain a bivariate normal distribution. In line with previous studies (53), the coefficients were then turned into absolute values and averaged across the moving windows.

#### 4.7.2. Expert raters judging the Solo session

The video and audio recordings associated with the Solo session (2 minutes) were presented to a group of six postgraduate students majoring in musicology [all blind to the experiment’s purposes, all having at least 8 years of music experience (mean ± SD = 10.50 ± 3.39 years)]. These music students, as expert raters, were asked to evaluate how well the music pieces had been acquired by providing subjective ratings on the 7-point Likert scales. The ratings consisted of the following six aspects [adapted from (4)]: (*i*) *Intonation*: Pitch accuracy; (*ii*) *Melody*: Ability to accurately express the linear succession of musical tones; (*iii*) *Rhythm*: Effective expression of the timing of musical sounds and silences that occur over time. (*iv*) *Lyric*: Accuracy in singing the lyrics; (*v*) *Emotion*: Ability to effectively express the emotion of the song. Signs of high ability include emotional facial and vocal expression; (*vi*) *Overall Performance*: The overall ability to perform the music song. We averaged the ratings provided by different raters and confirmed that the final score had very high inter-rater reliability (intra-class correlation on six aspects = 0.704 to 0.960). Also note that the sum of the first five aspects (i.e., Intonation + Melody + Rhythm + Lyrics + Emotion) was perfectly correlated with the last one (i.e., Overall Performance), *r* = 0.97, *P* < 0.001.

### 4.8. Statistical tests

A 2 × 3 mixed-design Analysis of Variance (ANOVA) was used to analyze all results across conditions. This ANOVA included a within-dyad factor RELATIVE-PHASE (in-phase, anti-phase, and sham) and a between-dyad factor FREQUENCY (6 Hz and 10 Hz).

When analyzing the cross-correlation coefficients, we conducted one ANOVA for each of the 251 time lags. The results from this series of ANOVAs required correction for multiple comparisons. To do so, we used a cluster-based permutation test (54). Specifically, adjacent time lags associated with a significant alpha level (*P* < 0.05) were grouped into a cluster. Then, a cluster-level statistic was calculated by taking the sum of the *F*-values within the observed clusters. The largest cluster was retained. To evaluate the significance of such largest cluster, we further created permuted data by shuffling dyads’ averages, i.e. randomly assigning dyads’ data to (*i*) either 6 Hz or 10 Hz groups and, within dyads, and to (*ii*) RELATIVE-PHASE conditions. The significance level was assessed by comparing the cluster statistics from the original data with 1000 renditions of permuted data using the Monte Carlo method (thresholded at *P* < 0.05).

Expert ratings were analyzed using the same mixed-design ANOVA, one for each of the six aspects being evaluated. Significant (*P* < 0.05) main effects or interactions were followed by pairwise comparisons. Bonferroni correction was used to account for *post hoc* multiple comparisons within each aspect of expert ratings.

Finally, we conducted correlation and mediation analyses to explore potential relationships between distinct dependent variables: interpersonal movement synchrony and expert ratings. In particular, following up on the results from the previous analyses, we computed “Δ interpersonal movement synchrony”, indexing the relative increase of interpersonal synchrony in the in-phase vs. sham stimulation condition, as well as “Δ intonation learning performance”, indexing the relative enhancement of intonation performance following in-phase vs. sham stimulation condition. Next, we conducted a Pearson correlation between these measures, as well as a mediation analysis, using INDIRECT macro (55) implemented in SPSS (version 18.0, SPSS Inc., Chicago, IL, USA). For the mediation analysis, we entered frequency of the dual brain stimulation (6 Hz vs. 10 Hz) as the independent variable, Δ interpersonal movement synchrony as the mediator, and Δ intonation learning performance as dependent variable. The bootstrapping method embedded in INDIRECT was used to determine whether the mediation effect was different from zero with 95% confidence. The number of bootstrap samples was set to 5000. Confidence intervals for indirect effect were bias corrected (55). Data are available at https://osf.io/bqsd5/.

## Acknowledgements

We would like to thank Philippe Peigneux, Charline Urbain, Pierre Besson, and Yixuan Ku for their valuable and insightful comments on earlier drafts, Soterix Medical Inc. for their technical assistance in the implementation of dual brain stimulation, Tianchi Luan, Xiushuang Zhang, Xuan Huang, Xueying Wang, Xinyuan Xu, Shuhan Jia for their help in acoustic data organization. This work was supported by the National Natural Science Foundation of China (31872783), the China Scholarship Council (201706140082), and the outstanding doctoral dissertation cultivation plan of East China Normal University (YB2016011).

## Author contributions

Y. P., G. N., B. S., and Y. H. designed research; Y. P. and Y. Z. performed research; Y. P. analyzed data; and Y. P., G. N., B. S., Y. Z., and Y. H. wrote the paper.

## Conflict of interest

The authors declare no conflict of interest.

