## Supporting Information Appendix for "Improving Social Interactive Learning through Dual Brain Stimulation"

**Supplementary Results**

**Brain stimulation questionnaires.** In a subset of participants (n = 12), we tested whether participants could detect the brain stimulation. Results were not consistent with this hypothesis. When asked whether they could or could not perceive the stimulation following each block, participants were at chance level (52.8%), indicating that participants could not reliably distinguish sham vs. actual stimulation.

Mixed-design ANOVAs also assessed that side-effects were comparable across conditions (*F*s < 1.14, *p*s > 0.12). A subset of participants (n = 3) reported perceiving phosphenes (none of them were paired together).

**Table S1.** Counterbalancement of different stimulation frequencies (6 Hz and 10 Hz), relative-phases (in-phase, anti-phase, and sham), and learning contents (songs) across experimental blocks and dyads.

| ***Dyad #*** | ***Block 1*** | ***Block 2*** | ***Block 3*** |
| --- | --- | --- | --- |
| 1 | 6 Hz / in-phase / song 1 | 6 Hz / anti-phase/ song 2 | 6 Hz / sham / song 3 |
| 2 | 6 Hz/ in-phase / song 2 | 6 Hz / sham / song 1 | 6 Hz / anti-phase / song 3 |
| 3 | 6 Hz / anti-phase / song 1 | 6 Hz / in-phase / song 3 | 6 Hz / sham / song 2 |
| 4 | 6 Hz / anti-phase / song 2 | 6 Hz / sham / song 3 | 6 Hz / in-phase / song 1 |
| 5 | 6 Hz / sham / song 1 | 6 Hz / anti-phase / song 3 | 6 Hz / in-phase / song 2 |
| 6 | 6 Hz / sham / song 2 | 6 Hz / in-phase / song 3 | 6 Hz / anti-phase / song 1 |
| 7 | 6 Hz / in-phase / song 1 | 6 Hz / anti-phase/ song 2 | 6 Hz / sham / song 3 |
| 8 | 6 Hz/ in-phase / song 2 | 6 Hz / sham / song 1 | 6 Hz / anti-phase / song 3 |
| 9 | 6 Hz / anti-phase / song 1 | 6 Hz / in-phase / song 3 | 6 Hz / sham / song 2 |
| 10 | 6 Hz / anti-phase / song 2 | 6 Hz / sham / song 3 | 6 Hz / in-phase / song 1 |
| 11 | 6 Hz / sham / song 1 | 6 Hz / anti-phase / song 3 | 6 Hz / in-phase / song 2 |
| 12 | 6 Hz / sham / song 2 | 6 Hz / in-phase / song 3 | 6 Hz / anti-phase / song 1 |
| 13 | 10 Hz / in-phase / song 1 | 10 Hz / anti-phase/ song 2 | 10 Hz / sham / song 3 |
| 14 | 10 Hz/ in-phase / song 2 | 10 Hz / sham / song 1 | 10 Hz / anti-phase / song 3 |
| 15 | 10 Hz / anti-phase / song 1 | 10 Hz / in-phase / song 3 | 10 Hz / sham / song 2 |
| 16 | 10 Hz / anti-phase / song 2 | 10 Hz / sham / song 3 | 10 Hz / in-phase / song 1 |
| 17 | 10 Hz / sham / song 1 | 10 Hz / anti-phase / song 3 | 10 Hz / in-phase / song 2 |
| 18 | 10 Hz / sham / song 2 | 10 Hz / in-phase / song 3 | 10 Hz / anti-phase / song 1 |
| 19 | 10 Hz / in-phase / song 1 | 10 Hz / anti-phase/ song 2 | 10 Hz / sham / song 3 |
| 20 | 10 Hz/ in-phase / song 2 | 10 Hz / sham / song 1 | 10 Hz / anti-phase / song 3 |
| 21 | 10 Hz / anti-phase / song 1 | 10 Hz / in-phase / song 3 | 10 Hz / sham / song 2 |
| 22 | 10 Hz / anti-phase / song 2 | 10 Hz / sham / song 3 | 10 Hz / in-phase / song 1 |
| 23 | 10 Hz / sham / song 1 | 10 Hz / anti-phase / song 3 | 10 Hz / in-phase / song 2 |
| 24 | 10 Hz / sham / song 2 | 10 Hz / in-phase / song 3 | 10 Hz / anti-phase / song 1 |
